# PhenoXtract: combining Large Language Model and Knowledge Graph embedding to extract phenotypes from clinical descriptions

**DOI:** 10.64898/2026.06.22.733382

**Authors:** Silvia Berardelli, Galadriel Brière, Benjamin Loire, Federica De Paoli, Andrea Gazzo, Ivan Limongelli, Paolo Magni, Susanna Zucca, Anaïs Baudot

**Affiliations:** Department of Electrical, Computer and Biomedical Engineering, University of Pavia, Pavia, Italy; enGenome s.r.l., Pavia, Italy; Aix Marseille Université, INSERM, MMG, Marseille, France; R&D Servier Paris-Saclay Institut, Gif-sur-Yvette, France; Unità di Biologia Computazionale e Bioinformatica, Fondazione Policlinico Universitario Agostino Gemelli IRCCS, Rome, Italy; CNRS, Marseille, France

## Abstract

**Motivation:** Standardized phenotypic descriptions are essential for accurate diagnosis, yet clinicians and researchers face challenges in manually extracting and mapping phenotypes from scientific literature or patient clinical records to the Human Phenotype Ontology. Recent advances in deep learning offer new opportunities for automation. We developed PhenoXtract, a novel phenotype extraction approach that combines Large Language Models and Knowledge Graph embedding. PhenoXtract is a multistep pipeline that takes clinical descriptions as input, extracts candidate phenotype entities using large language models, and maps them to terms from an enriched version of the Human Phenotype Ontology, processed as a knowledge graph.

**Results:** Evaluation against expert-curated ground-truth datasets show a recall of 0.70 and precision of 0.85 for PhenoXtract, demonstrating concordance with manually extracted phenotypes, with a computation time of 10-20 seconds for each text analyzed. Moreover, PhenoXtract surpasses rule-based and deep learning-based state-of-the-art tools in two out of the three ground-truth datasets evaluated. These results suggest that hybrid approaches combining Large Language Models and Knowledge Graph embeddings represent a promising direction for automated clinical phenotyping at scale.

## Introduction

Phenotypic information is captured across diverse sources and formats, spanning scientific literature, clinical reports, or electronic health records [1]. Indeed, journal articles and books describe patient phenotypes in research studies, case reports, and disease descriptions, while clinical documentation produced by physicians and geneticists includes detailed accounts of symptoms and conditions alongside laboratory, imaging, and genomic findings. In addition, clinicians routinely rely on free-text annotations to record and track patient phenotypes over time [2]. Standardized phenotypic descriptions are essential in healthcare because they reduce ambiguity in clinical interpretation and enable precise communication of patient features [3]. By using controlled vocabularies, phenotypes can be consistently captured and reliably interpreted by healthcare professionals across institutions and countries, facilitating the reuse of clinical findings without loss of meaning. Moreover, such standardization enables computational analysis of phenotypes across patients and diseases, supporting large-scale clinical research and data-driven discovery.

The Human Phenotype Ontology (HPO) is a structured, standardized ontology designed to describe phenotypic abnormalities observed in human diseases. By providing an unambiguous vocabulary for clinical symptoms, physical findings, and laboratory results, HPO facilitates consistent phenotypic data representation [4]. In recent years, the HPO has undergone regular updates, expanding its coverage and usability.

Despite these advantages, clinicians and researchers often struggle to manually extract and map phenotypic information from scientific literature or medical records into standardized HPO terms, a process that is time-consuming and prone to error [5]. Clinical descriptions, indeed, can vary widely in terms of wording and structure. Moreover, identifying and selecting the correct ontology terms is challenging, as it requires expert knowledge and constant navigation on large and complex terminologies. Consequently, there is a high demand for automated tools capable of efficiently mapping clinical text into standardized HPO terms [6].

Over the past years, several computational methods have been developed to automatically extract biomedical entities from clinical text, including phenotypic information mapped to standardized vocabularies. These approaches can be broadly categorized into rule-based, machine-learning and deep-learning based and LLMs-based methods [7].

Rule-based systems represent the earliest and most widely adopted category. They rely on dictionary matching and linguistic rules to identify phenotypes in text. Tools such as Doc2HPO [8] and ClinPhen [9] combine string matching with configurable options such as partial matching, negation detection, and context filtering to improve precision. More recently, FastHPOCR [10] introduced an optimized dictionary-based strategy designed to efficiently handle frequent HPO updates and large-scale corpora, emphasizing speed and ease of maintenance. Overall, rule-based approaches are transparent and easy to deploy, but their reliance on predefined rules limits their ability to capture complex linguistic context, often leading to false positives or false negatives results [11][12]. Additionally, rule-based systems require continuous refinement of the rules to cover evolving clinical terminology, limiting scalability.

Over recent years, machine-learning and deep-learning-based methods have been introduced, leveraging contextual language representations to better handle linguistic variability. Phenotagger [13], for example, combines a phenotype-oriented language model with dictionary matching to improve recognition beyond exact lexical overlap. PhenoBERT [14] further integrates ontology awareness into a neural pipeline, coupling candidate generation informed by the HPO hierarchy with a BERT-based evaluator. By explicitly modeling hierarchical relationships between phenotypes, PhenoBERT achieves improved accuracy, particularly in challenging cases where exact term matches are absent [14]. While these learning approaches offer substantial gains in performance, they depend on high-quality and large-scale training data and typically require retraining to adapt to ontology updates. More recently, LLM-based generative approaches have been explored for phenotype extraction. Studies investigating the use of general-purpose models such as ChatGPT highlight their ability to recognize phenotype mentions and map them to HPO terms, but also reveal significant limitations, including a lack of awareness of HPO updates, since it relies on pretrained knowledge [15]. Hybrids between LLM and rule based strategies, such as REAL [16], suggest that carefully engineered prompts and integration with knowledge sources as updated HPO terms details (synonyms, definitions) can partially mitigate these limitations. Nevertheless, LLM-driven solutions remain challenged by high computational costs, limited interpretability, and the need for frequent retraining or fine-tuning to accommodate the continuous evolution of the HPO [17]. These limitations motivate the development of methods that combine robust language modeling with explicit ontology knowledge, while remaining scalable to frequent updates in the HPO ontology.

We propose PhenoXtract, a hybrid method that combines the capabilities of LLMs with the structured, curated knowledge available in the HPO (Fig. 1). This design choice eliminates the need for retraining or fine-tuning the LLM when the ontology is updated, as newly introduced or modified HPO terms can be incorporated by updating the underlying ontology. PhenoXtract demonstrates promising initial performance, achieving stronger results than existing tools on two curated, clinically-oriented ground-truth datasets; however, on GSC+, a widely used benchmark designed for phenotype recognition and extraction in biomedical abstracts, its performance remains below that of FastHPOCR and PhenoBERT.

**Fig. 1.**
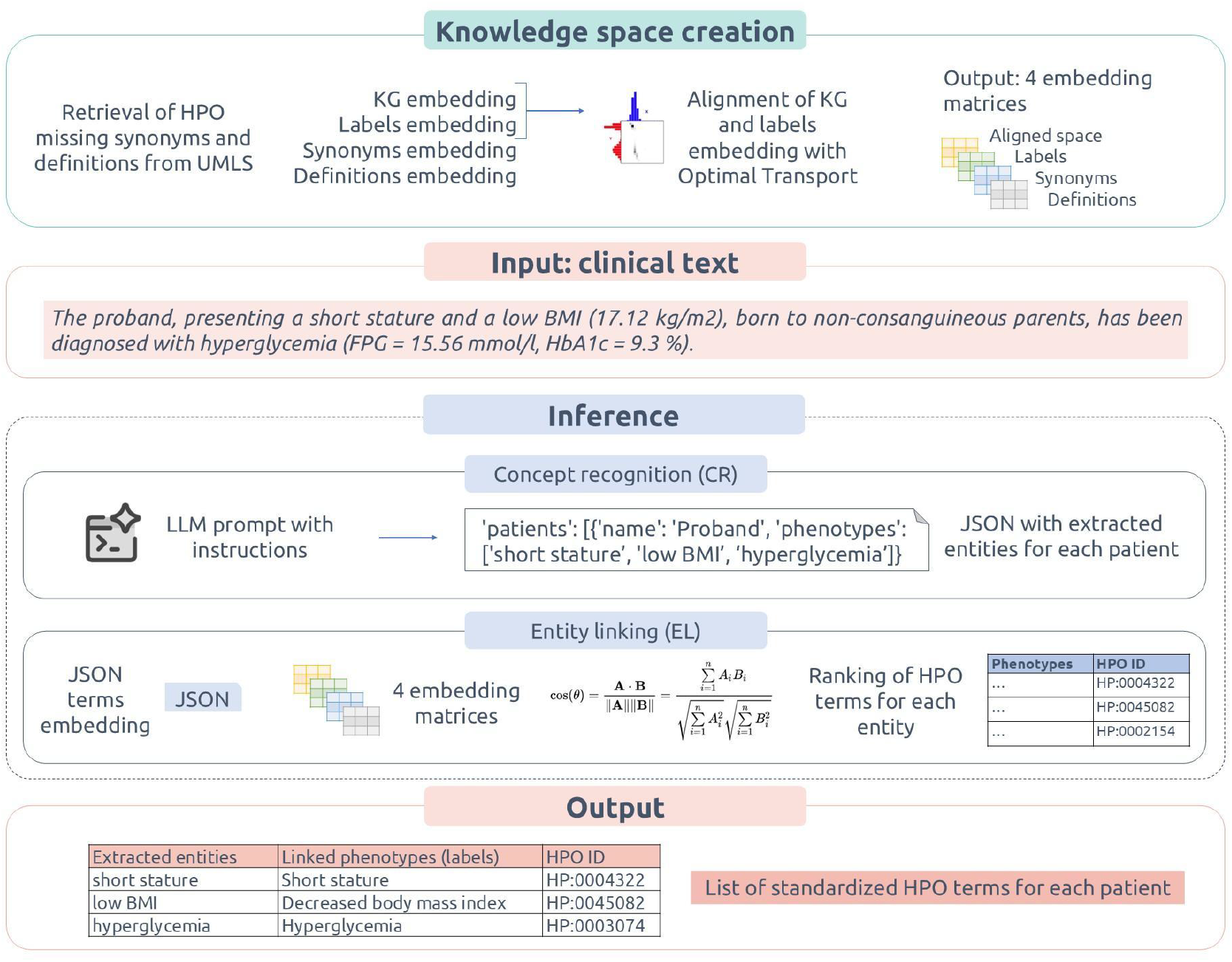
PhenoXtract workflow. Knowledge space creation involves the preprocessing of HPO. HPO terms lacking synonyms or definitions are retrieved from UMLS, when possible. HPO can be conceptualized as a Knowledge Graph (KG) consisting of nodes (phenotypic terms) and directed edges denoting hierarchical or relational links. We compute embeddings for the HPO KG, labels, synonyms, and definitions. We further aligned KG and labels embeddings to create the aligned space embedding. The input of the inference module for PhenoXtract is raw clinical text. Clinical text is processed with a two step approach: CR identifies and isolates entities; EL maps the entities to HPO terms, when possible. The output of PhenoXtract is a curated list of HPO terms corresponding to the input text.

## Material and Methods

Although phenotype descriptions can appear in a variety of sources, this work focuses on generic clinical text,including scientific literature and patient clinical reports, making our method source-agnostic.

PhenoXtract consists of an initial module of knowledge space creation, followed by a module for inference two steps: i) Concept Recognition (CR) and ii) Entity Linking (EL) (Fig. 1). CR is the task of automatically identifying and classifying entities within a text. Unlike simple keyword detection, CR focuses on higher-level semantic understanding of text, identifying underlying ideas or topics rather than just isolated words. In PhenoXtract, CR is applied to scientific literature and patient clinical records to extract entities that may be describing phenotypic features.

EL is the process of associating mentions of entities identified in text with corresponding unique entries in a knowledge base or database. EL involves disambiguation (distinguishing entities with similar names) and helps in creating structured representations of unstructured text data [18]. EL maps the entities extracted in the previous step of CR to HPO terms or determines if no mapping is possible.

### Knowledge space creation module: HPO preprocessing and embeddings creation

Each HPO term includes a primary label, a numerical identifier, a textual definition detailing the phenotype, associated synonyms, cross-references to external biomedical resources (e.g., OMIM [19], ORPHA [20], and SNOMED [21]), and hierarchical relationships within the ontology. The HPO hierarchical structure takes the form of a directed acyclic graph, where terms can have multiple parents, enabling accurate representation of complex biological relationships. Over recent years, the HPO has been regularly updated, resulting in a steady increase in both its coverage and usability (Supp. Fig. S1). Since 2021, curators added 2239 new terms and 49235 new disease-phenotype annotations, in collaboration with domain experts [4].

Knowledge space creation consists of three main steps: i) download of the HPO ontology in OWL format; ii) removal of the obsolete HPO terms from the ontology; iii) completion of the missing HPO terms definitions and synonyms, when possible. To accomplish step iii), we first perform a preprocessing step to address incomplete term annotations (examples provided in Supp. Fig. S2), such as missing synonyms or definitions. HPO terms with already present both synonyms and definitions require no modification, while incomplete entries are enriched using the UMLS API [22], which leverages cross-references across biomedical ontologies to retrieve missing information. After the removal of obsolete terms, HPO presents 18989 terms, where 8208 terms lack synonyms (43.2% of the total), and 2596 terms lack definitions (13.6% of the total). Thanks to our retrieval strategy, we retrieve 97% of missing synonyms and 99% of missing definitions.

Next, we adopt two representation strategies: (i) a textual embedding strategy based on term descriptions (for term labels, synonyms, definitions) and (ii) a KG-based strategy that leverages the HPO graph structure. For the textual embedding strategy, to obtain the three independent embedding matrices of textual data (label embedding, synonym embedding, definition embedding), the all-MiniLM-L6-v2 model from Sentence Transformers is applied (Supp. Table S4), as this model is recognized for its efficiency and reliable performance in large-scale contexts [23]. All enriched HPO terms labels, synonyms, and definitions are embedded across 384 dimensions. For the KG-based strategy, we conceptualize HPO as a KG consisting of nodes (HPO terms) and directed edges denoting hierarchical links [24]. We create the embedding with PyTorch Geometric [25] GraphSAGE package [26]. KG embedding enables the hierarchical organization and cross-term dependencies of HPO to be reflected in the learned representations.

We obtain four matrices: label embedding, synonym embedding, definition embedding, and KG embedding. Since textual embeddings and KG embeddings have been obtained with different models, aligning them into a unified space is essential. To achieve the alignment between label embedding and KG embedding, we employ Optimal Transport (OT) [27] (Supp. S.1.1). The final result, a unified embedding representation, will be referred to as aligned space embedding. As a final output of the knowledge space creation module, we obtain four distinct embedding matrices: label embedding, synonym embedding, definition embedding, and aligned space embedding. PhenoXtract uses all of those embeddings in the inference module (Supp. Table S5).

### Input

The expected input consists of clinical texts in English language for one or multiple patients (Supp. Fig. S3 for example). PhenoXtract accepts raw text input directly in string format. Alternatively, to facilitate the usage of PhenoXtract, the abstract of a scientific paper identified by a PMID or PMC identifier can be provided as input. In this case, we implemented an automated retrieval process: starting from the PMID or PMC identifier, a dedicated API call is triggered to automatically fetch the content of the abstract in string format or even the paper full text, whenever it is available [28].

### Inference module: Concept Recognition (CR)

In the CR step, a LLM-based approach is prompted explicitly to analyze clinical texts. Two different models are used, both provided by OpenAI: gpt-4o [29] and o3-mini [30]. Generic models, such as gpt-4o, are designed for broad tasks such as conversation, content generation, and multi-modal input processing. Reasoning models, such as o3-mini, are optimized for tasks requiring logic, problem solving, and step-by-step decompositions [31]. For the gpt-4o, the temperature parameter is set to 0.2. Temperature is a parameter that controls the randomness and creativity of the generated text by adjusting the probability of the next word [32]. The 0.2 value aims to reach a compromise between avoiding hallucinations and allowing the LLM to exploit context and its knowledge to detect phenotypic descriptions. In reasoning models, temperature setting is not allowed. The LLM is instructed with a structured zero-shot prompting methodology including four sections, for both gpt-4o and o3-mini (Supp. S.1.2). The Role and Instructions explicitly guides the model to perform CR through text segmentation and to distinguish between positive and negative phenotypic manifestations. In clinical terms, a positive phenotype corresponds to a trait explicitly observed in a patient. Conversely, negative refers to a trait explicitly stated as absent or ruled out in a patient; such information can be critical for differential diagnosis. When processing texts describing multiple patients, the prompt guides the LLM in stratifying phenotypic information at the level of individual patients or patient cohorts to ensure that annotations are not confused between individuals. The Input Text section includes raw, unformatted clinical text. The input text is segmented in predefined-length chunks to ensure that each chunk remains within the maximum token limit supported by the LLM, while still preserving sufficient contextual information for accurate phenotype extraction. The Safety Measures section guides the model in mitigating speculation and bias in phenotype identification. Finally, we instruct LLM to formulate the resulting Structured Output in JSON format, with each patient’s data encapsulated into discrete records containing the patient’s name or qualitative description, and the lists of explicitly identified positive phenotypes and negative phenotypes, respectively. The structured JSON output is intended as input for the subsequent steps. In the following steps, the phenotypes (both positives and negatives) extracted in the CR step will be referred to as entities.

### Inference module: Entity Linking (EL)

#### EL involves three main substeps

i. textual embedding of each entity extracted by CR using the all-MiniLM-L6-v2 model (Fig. S4A);
ii. comparison of the embedding of the extracted entity with the embedding matrices of the knowledge space different strategy for the comparison with the three textual embedding matrices (labels, synonyms, definitions) and the aligned-space embedding matrix. The comparison of the extracted entity embedding with textual embedding matrices (labels, synonyms, definitions) is performed directly through cosine similarity calculation and ranking (Supp. Fig. S4B); The comparison of the extracted entity embedding with the aligned space embedding involves first the processing of the extracted entity embedding vector with:
  a. retrieval of the 10 HPO terms most similar to the embedded entity (based on cosine similarity) in the label embedding matrix (as described in Eq. 1)

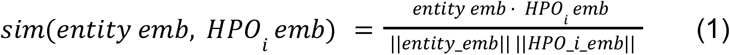
  b. weighted linear combination of the embedding vectors of these 10 HPO terms, with weights equal to cosine similarities, in order to approximate the entity’s position in the aligned space by anchoring it to its closest HPO neighbours. The ii)b substep allows the entity embedding to inherit the alignment properties of the label embedding and enable similarity comparisons within the aligned space. The output of the substep ii) is a new vector with the same embedding dimension as the label embedding dimension produced with textual embedding.
iii. Each entity is finally compared to HPO terms using cosine similarity across the four embedding matrices: labels, synonyms, definitions, and aligned space. Textual embedding is used for the first three comparisons, while weighted linear combination is used for the aligned space comparison. A link to an HPO term is established only when the similarity is significant as measured by cosine similarity.

The outputs of EL are lists of ranked HPO terms for each extracted entity and for each embedding category (label, synonym, definition, and aligned space). We save four top-ranking HPO terms, taking the one top-ranked from each embedding category for a final step of HPO terms linking. The aim of the final step is to determine which of the four top-ranked HPO terms from each embedding category should be selected. The implemented strategy follows different rules (Supp. Table S6), based on the level of agreement obtained with the different 4 top-ranked terms obtained from the 4 embeddings categories (examples in Supp. Table S1). We consider a candidate entity as an ambiguous case if it shows disagreement between the 4 top-ranked terms derived from the 4 different embedding categories. We implemented an LLM-based disambiguation step in these cases (example in Supp. Table S2). The final output of EL consists of a curated list of HPO terms, each including the original entity exactly as it appears in the input text, the corresponding standardized entity mapped to the HPO, and the associated unique HPO identifier (Supp. S.1.3).

### Ground-truth datasets and Evaluations

For the evaluation of the performances of PhenoXtract with both CR models (gpt-4o and o3-mini) and benchmarking comparisons with other approaches, we need manually curated ground-truth. We utilize 3 complementary ground-truth datasets, annotated by experts.

The curated datasets with a limited size enables detailed error analysis. The OLIDA (OLIgogenic Diseases DAtabase, https://olida.ibsquare.be/) is a database of digenic and oligogenic variant combinations associated with diseases [33]. It currently includes 1 808 oligogenic variant combinations linked to 219 distinct diseases. For each variant combination, OLIDA reports the PubMed identifiers (PMIDs) of the publications describing the association. From OLIDA, we selected a subset of 20 publications (Table S7). To build the PhenoXtract ground truth, a domain expert manually curated phenotypic annotations for each OLIDA entry by reviewing the associated publications. Clinical phenotypes were annotated by the expert using HPO terms and labeled as either positive or negative phenotypes. On average, the expert extracted 9 HPO terms for each OLIDA publication, focused on one disease, describing one or more patients.

The Mitochondrial ground-truth dataset, created internally in enGenome in 2023, includes clinical descriptions of 35 patients or cohorts of patients with mitochondrial pathogenic variants [34]. Each clinical description is associated with a publication record. The same expert curated the dataset, defining both positive and negative phenotypes associated with each patient, for an average count of 11 HPO terms. Because mitochondrial diseases typically present with complex phenotypes [35], this dataset is well suited to stress-test PhenoXtract under realistic diagnostic complexity.

The HPO Gold Standardized Corpora (GSC+) ground-truth represents a set of annotated biomedical texts optimized for testing computational methods [11][36]. It consists of 228 manually annotated abstracts, containing a total of 1933 HPO terms, with an average count of 9 HPO terms for each abstract, covering 460 unique HPO entities.

The three ground-truth datasets differ in several characteristics that are relevant for interpreting the evaluation results. The OLIDA and mitochondrial datasets were curated internally to capture clinically meaningful phenotype profiles from disease- and patient-oriented publication records. In both cases, expert annotation was performed at the level of the publication entry or patient/cohort description, with phenotypes categorized as positive or negative with respect to the reported condition. By contrast, GSC+ was designed as a benchmark corpus for phenotype CR and EL in biomedical text and consists of manually annotated PubMed abstracts with identified HPO annotations. GSC+ presents concise abstracts, with lexical variation and limited context.

Our aim is to evaluate the accuracy of PhenoXtract, with both the models, gpt-4o and o3-mini, in identifying positive and negative phenotypic descriptions in clinical text by comparing its annotations against manual, expert-curated ground-truth. In addition, PhenoXtract is benchmarked against two State-Of-The-Art (SOTA) phenotype extraction methods: the deep learning-based approach PhenoBERT, and the rule-based approach FastHPOCR. PhenoBERT is a deep learning framework that leverages contextualized language models to perform semantic similarity and normalization of clinical phenotype descriptions relative to the HPO. FastHPOCR is a dictionary-based concept recognition pipeline combining efficient boundary detection (i.e., the identification of the exact start and end of each entity in text) with entity linking. Its core idea is to reduce lexical variability by mapping entities to a large, pre-built set of clusters of morphologically equivalent entities. It emphasizes computational speed and scalability while maintaining high precision, making it suitable for large-scale biomedical text mining applications. For the evaluation, we employ precision, the proportion of correctly identified entities out of all entities predicted by our approach and benchmarked tools; recall: the proportion of correctly identified phenotypes or entities out of all actual phenotypes or entities annotated in the ground-truth; F1 Score, the harmonic mean of precision and recall, providing a balanced measure that takes into account both false positives and false negatives. In addition, we consider execution time for both CR and EL steps.

### Implementation and availability

PhenoXtract is a command-line tool and accepts as input parameters the input file with the text to be analyzed, the LLM model to be used, and the path to the output files.

We developed PhenoXtract entirely using Python 3.9. Scripts are available at: https://github.com/SilviaBerardelli/PhenoXtract.git

## Results

### Implementation of PhenoXtract

We developed PhenoXtract, which combines Large Language Models (LLMs) and Knowledge Graph (KG) embedding in a two-steps approach. In Concept Recognition (CR), PhenoXtract identifies candidate entities in clinical texts and in Entity Linking (EL), it maps phenotypic descriptions from clinical text to Human Phenotype Ontology (HPO) terms. An example of complete case study extracted from OLIDA dataset with the input-output workflow is described in Supp. S.1.3, providing step-by-step outputs of the method.

### Evaluation on ground-truth datasets

For each ground-truth dataset (OLIDA, Mitochondrial dataset, GSC+), we evaluate the number of phenotypes extracted with PhenoXtract (Supp. Fig. S5). A median of 10 and 8 HPO terms per paper are extracted with gpt-4o and o3-mini, respectively, from the OLIDA dataset. PhenoBERT extracted a median of 10 HPO for the OLIDA dataset and FastHPOCR extracted a median of 9 HPO terms. A median of 15 and 12 HPO terms per paper are extracted with gpt-4o and o3-mini from the mitochondrial dataset. PhenoBERT extracted a median of 14 HPO for the OLIDA dataset and FastHPOCR extracted a median of 13 HPO terms. A median of 4 HPO terms per abstract are extracted with both gpt-4o and o3-mini from the GSC+ dataset. Both PhenoBERT and FastHPOCR extracted a median of 5 HPO for the GSC+ dataset. Overall, the two models, gpt-4o and o3-mini, extract comparable numbers of phenotypes.

On the OLIDA ground-truth (Fig. 2A, Table 1), PhenoXtract with gpt-4o model (F1-score=0.78±0.19, mean±sd) and PhenoXtract with o3-mini model (F1-score=0.76±0.12) outperforms the two other methods from the SOTA across all three metrics (PhenoBERT F1-score=0.57±0.19, FastHPOCR F1-score=0.52±0.16), showing higher median and mean recall and precision, and consequently higher F1-scores. PhenoXtract with gpt-4o achieves performance with narrow inter-quartile ranges and less outliers, whereas o3-mini performs slightly lower but still superior to the SOTA systems.

**Table 1.** Benchmark results. PhenoXtract results compared to PhenoBERT and FastHPOCR on the three ground-truth datasets. Results are reported with median/mean (sd) format. Best results are in bold.

| Ground-Truth | Model | Recall | Precision | F1 |
| --- | --- | --- | --- | --- |
| OLIDA | PhenoXtract (*gpt-4o*) | **0.77/0.75** (0.22) | **0.90/0.84** (0.22) | **0.82/0.78** (0.19) |
| OLIDA | PhenoXtract (*o3-mini*) | 0.71/0.72 (0.19) | 0.89/**0.85** (0.12) | 0.78/0.76 (0.12) |
| OLIDA | FastHPOCR | 0.50/0.52 (0.15) | 0.62/0.55 (0.18) | 0.57/0.52 (0.16) |
| OLIDA | PhenoBERT | 0.62/0.56 (0.20) | 0.67/0.61 (0.23) | 0.64/0.57 (0.19) |
| Mitochondrial | PhenoXtract (*gpt-4o*) | **0.87/0.86** (0.12) | **0.88/0.85** (0.11) | **0.86/0.85** (0.10) |
| Mitochondrial | PhenoXtract (*o3-mini*) | 0.85/0.85 (0.13) | 0.85/0.84 (0.12) | 0.84/0.84 (0.11) |
| Mitochondrial | FastHPOCR | 0.56/0.57 (0.20) | 0.56/0.53 (0.19) | 0.56/0.54 (0.17) |
| Mitochondrial | PhenoBERT | 0.56/0.56 (0.21) | 0.58/0.58 (0.21) | 0.58/0.56 (0.19) |
| GSC+ | PhenoXtract (*gpt-4o*) | 0.36/0.37 (0.24) | 0.67/0.58 (0.32) | 0.44/0.42 (0.23) |
| GSC+ | PhenoXtract (*o3-mini*) | 0.33/0.36 (0.25) | 0.59/0.58 (0.32) | 0.44/0.42 (0.23) |
| GSC+ | FastHPOCR | **0.50/0.49** (0.26) | 0.67/0.62 (0.30) | 0.56/0.53 (0.25) |
| GSC+ | PhenoBERT | 0.50/0.47 (0.25) | **0.75/0.72** (0.30) | **0.57/0.55** (0.24) |

**Fig. 2.**
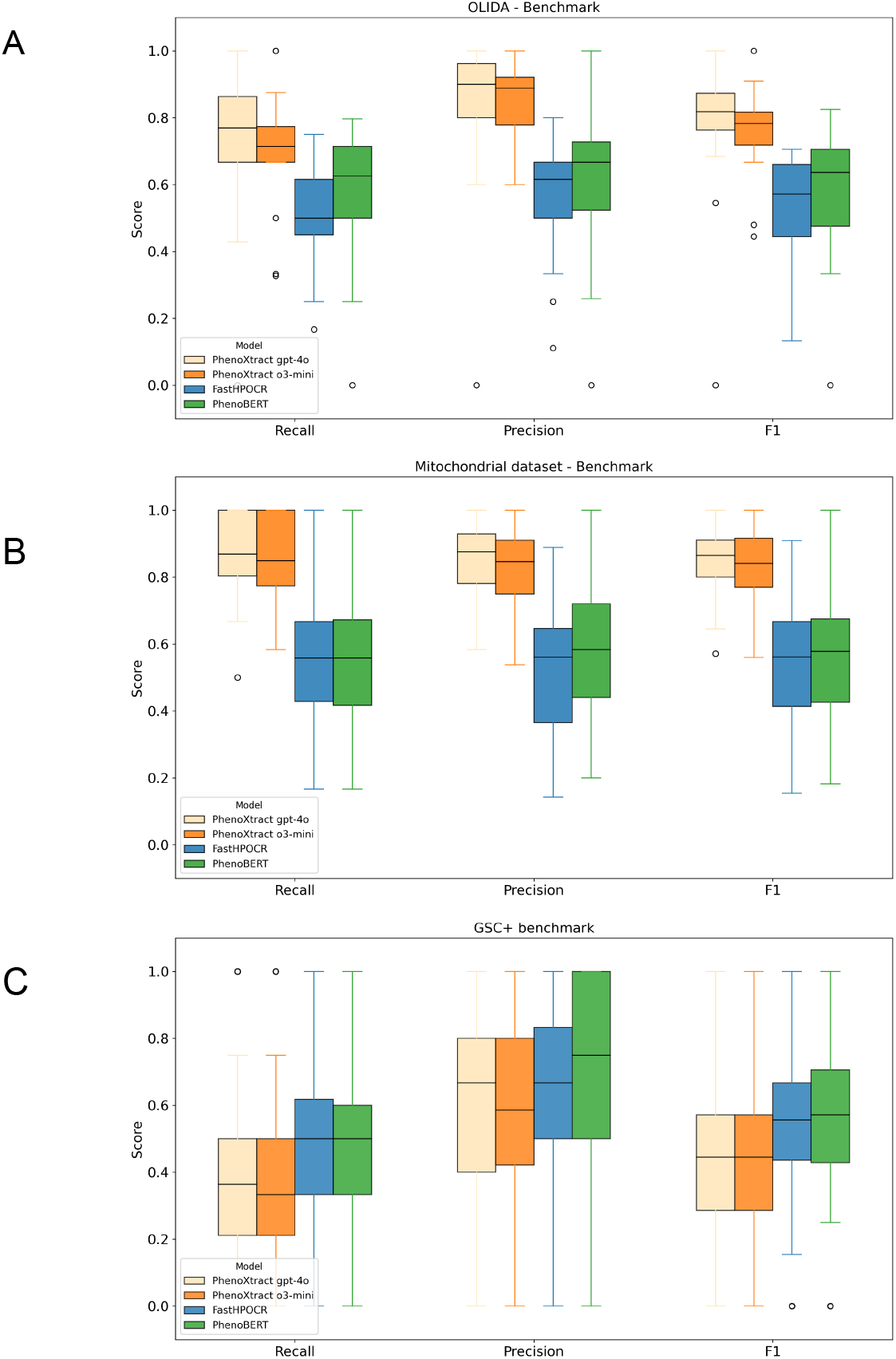
PhenoXtract performance comparison against PhenoBERT and FastHPOCR. Benchmark boxplots for OLIDA (2A), Mitochondrial (2B) and GSC+ (2C) datasets.

On the mitochondrial disease benchmark (Fig. 2B), the PhenoXtract with gpt-4o achieves the highest performance, with a median recall of 0.87 (0.86±0.12, mean±sd), precision of 0.88 (0.85±0.11), and F1-score of 0.86 (0.78±0.1), with low variability. The PhenoXtract with o3-mini implementation performs comparably, with lower but still robust median values (recall 0.85, precision 0.85, F1-score 0.84). In contrast, FastHPOCR and PhenoBERT demonstrate lower performance, with median F1-score of 0.56 and 0.58, respectively, and wider variability across clinical texts.

When evaluated on the GSC+ benchmark (Fig. 2C), both FastHPOCR and PhenoBERT outperform PhenoXtract, reaching median F1-scores of 0.56 and 0.57, respectively. The precision of PhenoBERT (median 0.75) and FastHPOCR (median 0.67) compensates for their moderate recall, whereas PhenoXtract shows lower recall (median 0.36). All tools display lower performance compared to the OLIDA and mitochondrial datasets. PhenoXtract, implemented with gpt-4o and o3-mini, achieves median F1-scores of 0.44, substantially lower than those observed on OLIDA (0.82 and 0.78, respectively) and mitochondrial disease (0.86 and 0.84, respectively). GSC+ appears to be the most challenging dataset for PhenoXtract (see examples and additional tests in Supp. S.1.4).

Overall, on the OLIDA and mitochondrial datasets, which provide curated, disease specific corpora, PhenoXtract (particularly the gpt-4o implementation) achieves the highest performance, with F1-scores exceeding 0.80 and stable variance across clinical texts. FastHPOCR and PhenoBERT exhibit broader variability and lower median scores, especially for recall and F1, indicating less robust extraction of HPO terms from the benchmark corpus. The comparison between PhenoBERT and FastHPOCR shows broadly comparable performance, with PhenoBERT exhibiting a slight tendency toward higher median precision and F1 scores, while maintaining a similar level of variability across the evaluated benchmarks.

In contrast, performance on the GSC+ is substantially lower, with median F1-score around 0.44 for PhenoXtract. PhenoBERT and FastHPOCR achieved comparatively better results, particularly due to higher precision, suggesting that PhenoXtract struggles to generalize when applied to more variable biomedical text.

### Processing times evaluation

The observed differences in PhenoXtract processing times across datasets (Supp. Fig. S6, and Supp. Table S8) are primarily explained by the length of the clinical input texts (Supp. Fig. S7), which is substantially greater for OLIDA and the mitochondrial dataset compared to GSC+, because GSC+ is based only on abstracts. Moreover, across all datasets, substantial differences were observed in total execution time comparing PhenoXtract to SOTA. PhenoXtract, relying on LLM inference through API calls, consistently exhibits the highest runtimes. In contrast, PhenoBERT and FastHPOCR approaches show markedly lower total processing times and compact distributions, highlighting a clear separation between API-dependent pipelines and non-LLM approaches in terms of computational efficiency.

Since the goal is to build a versatile and robust approach that could also be used with other ontologies, we conduct initial tests of the approach with the Disease Ontology (DO), which classifies rare and common human diseases [37]. We apply our two-steps CR/EL approach to recognize and extract disease descriptions and map them to standardized DO terms, which only requires adaptations involving the ontology file and the LLM prompt used in the CR step. Initial examples show promising results (Supp. Fig. S8).

## Discussion and future directions

While LLMs have rapidly become a widespread technology and have allowed the development of a plenty of tools for clinical text processing, their exclusive use still presents limitations. One of the main challenges lies in the need for retraining and fine-tuning, which makes relying solely on LLMs often ineffective when the goal is to achieve domain-specific up-to-date terminology extraction. In this sense, PhenoXtract represents a hybrid strategy combining the strengths of LLMs and KG embedding for HPO terms extraction, scalable to be applied to other ontologies. By leveraging the HPO structure, PhenoXtract ensures that the identified HPO terms are not only accurate but also consistently mapped to standardized entries. Moreover, the integration with UMLS enriches the ontology coverage by providing synonymy and detailed descriptions, especially for HPO terms that are otherwise poorly represented. Within reasonable computational times, PhenoXtract transforms unstructured text or tabular clinical data into a curated list of HPO terms with their corresponding identifiers, offering clinicians and researchers a reliable representation of patient phenotypes.

Presented results demonstrate that PhenoXtract achieves very good performance on curated corpora such as OLIDA and mitochondrial datasets, consistently outperforming existing deep learning-based and rule-based tools. The evaluation on theGSC+ benchmark reveals important limitations of PhenoXtract, including sensitivity to prompt design choices, issues with ontology versioning (see examples and additional tests in Supp. S.1.4), and reduced generalizability in heterogeneous text corpora.

A challenge in the proposed pipeline and for all the LLMs-based approaches concerns the intrinsic lack of reproducibility of steps relying on LLMs [38]. LLM-based tasks inherently depend on probabilistic generation, which may lead to variability across runs even when identical inputs are provided. This stochastic behavior limits strict reproducibility and complicates the validation of results in sensitive biomedical contexts. In [39], the authors propose a statistical framework for evaluating LLM consistency by quantifying the reproducibility of outputs in diagnostic reasoning. The authors highlight the need for case-by-case assessment of output consistency to ensure LLMs reliability in clinical applications.

The execution time of the CR step is not scalable, as the text is sent to the LLM via a single call. One possible improvement would be to split the text into smaller chunks and launch parallel calls, but this would also increase costs. The adoption of LLMs in clinical applications entails significant costs, including expenses related to inference, infrastructure, and data privacy compliance, which may limit scalability in real-world settings. To mitigate these challenges, deploying optimized or smaller models locally can represent a viable alternative.

For these reasons, we recommend its use in settings where eventual careful review of extracted phenotypes is feasible, rather than as a replacement for highly optimized mention-detection tools in large-scale mining workflows.

A useful extension for PhenoXtract would involve the automated extraction of quantitative phenotypes from clinical records. These may appear in different forms: numerical values explicitly reported in structured tables and associated with individual patients; textual descriptions that encode quantitative information (e.g., systolic pressure over 150 mmHg); or qualitative statements referring to ranges or relative levels (e.g., high level of glucose in blood, above the normal threshold). The processing of such information requires handling challenges such as acronyms, measurement units, and variable reporting styles [40]. To support interoperability and semantic clarity, ontologies (such as UMLS, LOINC, SNOMED) can be used to enrich textual descriptions and map qualitative terms to standardized entities.

Beyond the HPO, a step for expanding the automated extraction is the integration of multiple biomedical ontologies into a unique comprehensive KG. This extension would allow the recognition and linking of diverse entity types, including genes, diseases, genetic variants, and drugs, thereby supporting a richer and interconnected representation of clinical and molecular information [41][42].

## Acknowledgements

The aims of this study contribute to the ERDERA project, which has received funding from the European Union’s Horizon Europe research and innovation programme under grant agreement N°101156595. This work was supported by the The French Muscular Dystrophy Association (AFM-Telethon).

