## Supplementary Material for "PhenoXtract: combining Large Language Model and Knowledge Graph embedding to extract phenotypes from clinical descriptions"

S.1. Additional Methodological Details

S.1.1 Optimal Transport

Optimal Transport provides a way to compare probability distributions by minimizing the total “cost” of transportation. The cost is typically defined in terms of distance, such as how far mass needs to be moved. Optimal transport has become a powerful tool across biology and bioinformatics, as described in [1], [2] and [3]. Its central object, the Wasserstein distance, captures both geometric and probabilistic differences between distributions [4]. Figure below shows examples of OT application in both 1D (Figures A, Figure B) and 2D contexts (Figure C, Figure D). In 1D context, two probability mass functions are defined on the interval [0, 1]. The cost matrix C along with the same density functions (one up top and other flipped vertically). The cost is zero along the diagonal of C since it costs us nothing to move mass zero units of distance. Defining the transportation cost as squared Euclidean distance, moving vertically or horizontally off the diagonal increases the cost quadratically. The two target marginal distributions p and q and the cost matrix C produces the optimal transport plan T*, a matrix the same size as C. In 2D, discretization in spatial bins is performed. The 2D densities are flattened into 1D vectors. Since it is a 20 x 20 discrete grid, there are a total of 400 bins, and thus a 400 x 400 matrix. The 2D problem is reduced to the 1D, and the same idea to identify the optimal transport plan can be used. In figure below E, , the arrow schematizes units being transported from location (x0, y0) to (x1, y1). A complete transport plan specifies transport paths over all pairs of locations. Finally, figure below F shows a simplified visualization of the optimal transport plan for the (discretized) 2D example problem.

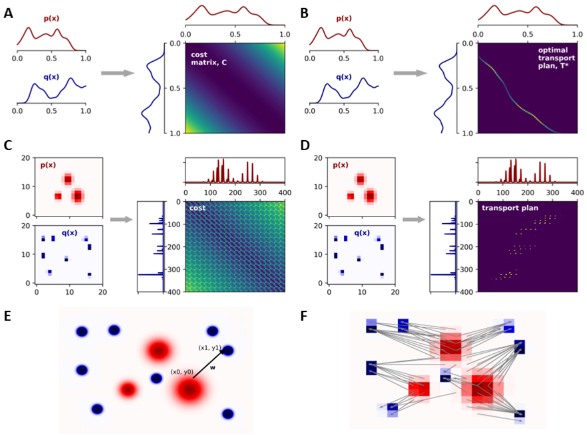


**Optimal transport plan matrix.** A: density functions for two 1D distributions for P and Q and cost matrix C showing squared Euclidean distances between all pairs of points. B: transport plan matrix T*. Entry (i, j) in this matrix specifies how much mass in bin i of Q should be transported to bin j of P . C: symmetric matrix of transport costs. The blocky structure arises because of the flattened 2D grid of bins. D: transport plan matrix T*. E: example of transport plan with transport paths for a pair of locations. F: optimal transport plan for a 2D example.

Applied to PhenoXtract, OT specifically targets alignment between label embeddings and KG embeddings due to their complementary semantic richness [5][6][7]. Python Optimal Transport (POT) package was exploited for this task. First, the cost matrix C is computed using the squared Euclidean distance between labels embeddings and graph embeddings (Eq. 2):

C = cdist(labels_embeddings, graph_embeddings, metric = 'seuclidean') (2)

This captures the pairwise costs of transporting “mass” from points in the text space to points in the graph space. Next, the Sinkhorn algorithm from the POT library is applied, where a and b are the distributions over the text and graph embeddings, and reg is the entropic regularization parameter that enforces smoothness in the solution. The resulting transport matrix G specifies how much mass from each point in the text space is assigned to each point in the graph space, with G[i, j] representing the amount transported from text point i to graph point j (Eq. 3). Finally, the aligned latent embeddings are obtained as a weighted combination of the graph embeddings (Eq. 4):

G = ot.sinkhorn(a, b, C, reg = 0.1) (3)

aligned_space_embeddings = G · graph_embeddings (4)

S.1.2 Concept Recognition LLM prompt

The complete prompt is reported:

***Role and Instructions**: Act as a clinical assistant: your aim is to support clinicians in the task of phenotype extraction from text. Your task is to produce a list of entities extracted from a clinical description, in the raw text provided, both for phenotypes and for negative phenotypes. Given an input text, you must identify the spans related to possible phenotypes, either explicitly or implicitly. You should keep in the span all words related to the phenotype that should be informative (such as negation or adjective). Into negative phenotypes, include also descriptions like “recovered without any neurological disability”, where it is specified that the patient does not have that symptom. Avoid keeping descriptions that are not mainly related to phenotypes, such as “male child” or “female newborn”. Avoid keeping descriptions related to mutations and mode of inheritance, as “mutational burden” or inheritance qualifiers. Avoid keeping descriptions as “asymptomatic”. Avoid keeping patients that have no phenotypes specified. You may reformulate the span if needed. If you recognize more than one patient in the text, please list*

*all the phenotypes owned by each patient. If you do not detect any span or if you do not know, do not try to make up an answer, just write ‘None’. If you detect phenotypes without patient specifications, use “not specified” for patients but include phenotypes.*
***Input text**: {content}.*
***Safety Measures**: Avoid redundancy. Avoid speculation: extract phenotypes known or explicitly available from the text. Comprehensive use of text: Utilize all relevant information across the entire text to ensure diversity.*
*Structured output:*


[{ "name": "extract_patient_phenotypes",
"description": "Extract phenotypes and negative phenotypes for
each patient from the text.",
"parameters": {"type": "object",
"properties": {
"patients": {
"type": "array",
"description": "List of patients with their phenotypes and
negative phenotypes.",
"items": {
"type": "object",
"properties": {
"name": {
"type": "string",
"description": "Name of the patient."},
"phenotypes": {"type": "array",
"items": {"type": "string"},
"description": "List of phenotypes for the patient."},
"negative_phenotypes": {
"type": "array",
"items": {"type": "string"},
"description": "List of negative phenotypes
for the patient."}},
"required": [
"name",
"phenotypes",
"negative_phenotypes" ]}}},
"required": ["patients" ]}}]


An example of structured output is reported:


{ "patients": [ {
"name": "30-year-old Chinese female patient",
"phenotypes": [
"chronic active Epstein-Barr virus infection",
"cutaneous lymphoproliferative disorders",
"haemophagocytic lymphohistiocytosis",
"fatigue",
"recurrent high-grade fever",
"uncontrolled splenomegaly",
"oedematous swelling of the cheeks, eyelids and lips",
"coexistent skin lesions",
"liver damage",
"pancytopenia",
"hypofibrinogenemia",
"macrophage activation syndrome",
"capillary leak syndrome",
"EBV infection"],
"negative_phenotypes": [
"abnormal lymphocytes or haemophagocytosis",
"no SH2DIA or XLP1 mutations found" ]}]


S.1.3 Case Study

The example reports a text extracted from OLI1146 patient (OLIDA dataset) [8].

*A Tunisian girl presenting a **short stature** and a **low BMI** (17.12 kg/m2), born to non-consanguineous parents, has been diagnosed with **hyperglycemia** (FPG = 15.56 mmol/l, HbA1c = 9.3%) at 21 years old after **recurrent signs of polyuria and polydipsia**. She started a healthy diet coupled with sulfonylurea uptake under the recommendation of her clinician. Besides diabetes, our patient has presented **intellectual disability** and **post-axial brachydactyly since birth**. Signs of **mental impairment** have remained during her childhood, with **speech disorders** and **learning difficulties**. Fourteen months later, the clinician noticed a **persistent hyperglycemia** (FPG = 13.2 mmol/l, HbA1c = 8%) as well as a low level of C peptide (1.08* *ng/ml). Therefore, an insulin therapy has been prescribed.* ***Abdominal and pelvic ultrasounds were normal****. Although the patient had* ***slightly low creatinine levels*** *(41 µmol/l), she had normal 24-h urine albuminuria (15 mg/24 h), as well as a* ***normal estimated glomerular filtration rate*** *(eGFR = 136.9 ml/min).*

Phenotypes identified by PhenoXtract CR, both “positive” and “negative” are shown in bold.
The output JSON of CR step is the following:

*{"patients": [{*
*"name": "not specified",*
*"phenotypes": [*
*"short stature",*
*"low BMI (17.12 kg/m2)",*
*"hyperglycemia at 21 years",*
*"recurrent signs of polyuria and polydipsia",*
*"intellectual disability since birth",*
*"post-axial brachydactyly since birth",*
*"signs of mental impairment during childhood",*
*"speech disorders",*
*"learning difficulties",*
*"persistent hyperglycemia",*
*"slightly low creatinine levels"],*
*"negative_phenotypes": [*
*"normal abdominal and pelvic ultrasounds",*
*"normal 24-h urine albuminuria (15 mg/24 h)",*
*"normal estimated glomerular filtration rate (eGFR = 136.9 ml/min)"]}]}*


Examples of Cosine similarity calculation, ranking and final EL to HPO term is shown for short stature (Table S1) and for speech disorders (Table S2). *Short stature* results are an example of the easiest case for EL: 3 out of 4 top-ranked phenotypes are with a 1.0 cosine similarity and are perfect matches. Contrarily, for speech disorders the 4 top-ranked phenotypes are different from each other, so an additional LLM step is used. The prompt used for the additional LLM step is reported:

***Role and Instructions:*** *Act as a genomics assistant: your aim is to support geneticists in the task of phenotype extraction from text. Your task is to identify, given a list of terms as candidates, the most fitting one for the query given. Query is a description of a symptom or condition extracted from a text. List of candidates is a list of up to four terms. Of course, query and candidates can be different. Your aim is to identify the most similar, for a semantic reason. If you don’t detect any fitting term at all or if you don’t know, don’t try to make up an answer, just write ’None’.*
***Query:*** *{query}*
***List of possible phenotypes between you have to choose:*** *{content}.*
***Safety Measures:*** *Avoid speculation: extract phenotypes known or explicitly available from the list.*
***Structured output:***

*[{ "name": "identify_patient_phenotypes",*
*"description": "Most fitting term from the provided list.",*
*"parameters": {*
*"type": "object",*
*"properties": {*
*"top_1_candidate": {*
*"type": "string",*
*"description": "Most fitting term from the provided list."}},*
*"required": ["top_1_candidate"]}}]*


An example of structured output is reported:

*{"top_1_candidate": "Dysphonia"}*


The output of EL is reported in Table S3.

**Table S1.** *Short stature* 4 top-ranked phenotypes.

| **Embedding category** | **Top-1 HPO Term** | **Cosine** |
|---|---|---|
| labels | short stature | 1.0 |
| synonyms | proportionate short stature | 0.81 |
| definitions | short stature | 1.0 |
| aligned space | short stature | 1.0 |

| **Embedding** | **Top-1 HPO Term** | **Cosine** |
|---|---|---|

| category | | |
|---|---|---|
| labels | Abnormal speech discrimination | 0.72 |
| synonyms | Dysphonia | 0.74 |
| definitions | Deficit in grammar | 0.83 |
| aligned space | Abnormal speech volume | 0.90 |

**Table S2.** *Speech disorders* 4 top-ranked phenotypes. The additional LLM step identifies *{"top_1_candidate": "Dysphonia"}*, so Dysphonia is the given output for Speech disorder

| entity | neg | HPO | HPO ID |
|---|---|---|---|
| short stature | 0 | Short stature | HP:0004322 |
| low BMI (17.12 kg/m2) | 0 | Decreased body mass index | HP:0045082 |
| hyperglycemia at 21 years | 0 | Hyperglycemia | HP:0003074 |
| recurrent signs of polyuria | 0 | Polyuria | HP:0000103 |
| intellectual disability since birth | 0 | Intellectual disability | HP:0001249 |
| post-axial brachydactyly since birth | 0 | Brachydactyly | HP:0001156 |
| signs of mental impairment during childhood | 0 | Developmental regression | HP:0002376 |
| speech disorders | 0 | Dysphonia | HP:0001618 |
| learning difficulties | 0 | Specific learning disability | HP:0001328 |
| persistent hyperglycemia | 0 | Hyperglycemia | HP:0003074 |
| slightly low creatinine levels | 0 | Abnormal circulating creatinine concentration | HP:0012100 |
| normal abdominal and pelvic ultrasounds | 1 | Abnormality of the abdominal organs | HP:0002012 |
| normal 24-h urine albuminuria (15 mg/24 h) | 1 | Moderate albuminuria | HP:0012594 |
| normal estimated glomerular filtration rate (eGFR=136.9 ml/min) | 1 | Abnormal glomerular filtration rate | HP:0012212 |

**Table S3:** EL output for OLIDA OLI1146. The neg column stands if the phenotype is positive (0) or negative (1)

#### S.1.4 Additional evaluation on GSC+

A closer inspection of false negatives and false positives in the GSC+ benchmark reveals that part of PhenoXtract's lower performance can be attributed to prompt design choices. For instance in sample ID: 1683160, describing Angelman Syndrome (AS):

*DNA deletion studies using 5 DNA markers localized at 15q11-q12 were performed in 14 Angelman syndrome (AS) patients (9 sporadic and 5 familial cases). A one-copy density for one or more of the 5 loci was detected in 8 (57.1%) of the 14 patients. A deletion of only the D15S11 locus was detected in one sporadic patient, that involving only the D15S10 in 3 familial patients (sibs in a family), that spanning 3 loci (D15S11, D15S10, D15S12) in one sporadic patient, and that spanning 4 loci (D15S9, D15S11, D15S10, D15S12) in the other 3 sporadic patients.*

In the sentence describing Angelman syndrome patients with sporadic and familial cases, the ground truth annotations include **Sporadic** (HP:0003745) and **Mode of inheritance** (HP:0000005). However, in the prompt there is this explicit instruction:

*Avoid keeping descriptions related to mutations and mode of inheritance, as mutational burden or inheritance qualifiers.*

Since it is explicit in the extraction prompt to avoid inheritance qualifiers, these phenotypic descriptors were excluded, resulting in missed annotations. At the same time, PhenoXtract extracted the term Angelman syndrome and mapped it to Abnormal nervous system electrophysiology (HP:0001311) (Figure below). PhenoXtract successfully identified Angelman syndrome (OMIM:105830), even though this entity was not part of the reference annotations 3. This example illustrates a dual source of error: on the one hand, excessively restrictive requests may prevent valid extractions, and on the other hand, the model may provide conceptually relevant but out-of-scope annotations. Both factors contribute to reduced recall on GSC+, particularly in cases where inheritance patterns or disease mentions are present in the text. Nonetheless, the fact that PhenoXtract captured a disease entity with established HPO-OMIM linkage suggests that even some "errors" may retain biological utility, underscoring the complexity of defining strict ground truth in heterogeneous biomedical corpora.

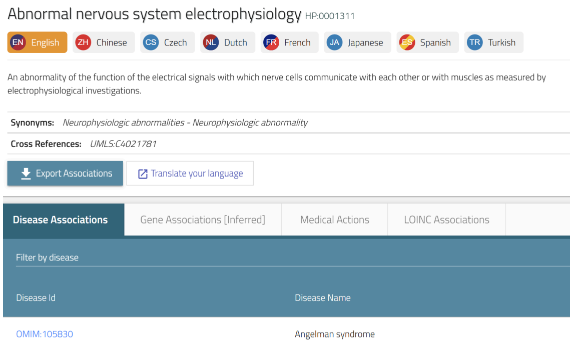


Abnormal nervous system electrophysiology term in HPO. The term *Abnormal nervous system electrophysiology* has Angelman syndrome between the Disease Associations.

Another source of apparent errors in the GSC+ arises from discrepancies between ontology versions, identified in sample ID: 3100017.

*In situ hybridization with a c-sis probe was performed on peripheral lymphocytes of a man with neurofibromatosis and a ring 22 chromosome. Hybridization was observed on both the normal #22 and the ring 22, indicating that the patient is not constitutionally hemizygous for c-sis. The implications of a ring 22 constitution and the neurofibromatosis phenotype are discussed.*

In the example sentence describing a patient with neurofibromatosis and a ring 22 chromosome, the ground truth annotation lists neurofibromatosis (HP:0006746), a term that has since been deprecated in the HPO. As a result, the automated comparison flagged this case as a false negative. However, PhenoXtract mapped the mention correctly to Neurofibroma (HP:0001067) (Figure below), which explicitly includes neurofibromatosis among its synonyms.

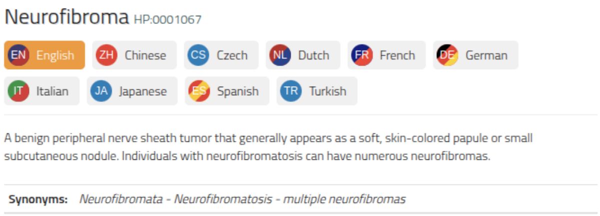


Neurofibroma term in HPO. The term Neurofibroma has neurofibromatosis between its synonyms. PhenoXtract correctly detects and maps it.

The example indicates that the extraction was a true positive, but it was penalized due to ontology versioning issues in the reference set. Such cases highlight the challenges of evaluating automated tools against evolving biomedical ontologies: what is counted as an error in strict automated matching may actually represent a biologically correct and up-to-date annotation. Careful consideration of ontology changes, and cross-version mapping is therefore essential to avoid underestimating model performance.

#### S.1.5 Reproducibility analysis of HPO extracted across repeated runs

Reproducibility analysis is a critical component in the evaluation of tools based on LLMs. In fact, LLMs may generate partially variable outputs across repeated executions, even when provided with the same input data. Thus, we conducted a quantification of the intra-sample stability of the generated outputs across independent runs. Given the cost of repeated LLM API calls, performing the reproducibility assessment on the full benchmark datasets was not computationally feasible. We therefore selected a subset of 10 samples from the OLIDA dataset and 10 samples from the Mitochondrial dataset. We repeated the extraction procedure 10 times for each sample. This setup provided a practical compromise between cost containment and the estimate of run-to-run variability of the extracted HPO terms.

For each sample, we computed pairwise Jaccard (Eq. S1) and Dice (Eq. S2) similarities between all run pairs and the proportion of terms consistently recovered across all runs, that we defined as the terms ratio T (Eq. S3).

$J(A,\, B) = \frac{|A \cap B|}{|A \cup B|}$ (S1) $DSC(A,\, B) = \frac{2|A \cap B|}{|A| \cup |B|}$ (S2) T = $\frac{\left|\bigcap_{r=1}^{10} s_{i,r}\right|}{\left|\bigcup_{r=1}^{10} s_{i,r}\right|}$ with $s_{i,r}$ extracted HPO terms for sample *i*, for run *r* (S3)

In the mitochondrial dataset, the mean pairwise Jaccard similarity is 0.907 and the mean Dice similarity is 0.947, indicating high agreement between repeated runs. The mean terms ratio was 0.761, suggesting that a large proportion of the HPO terms observed across runs belonged to a stable set extracted for each sample.

The OLIDA dataset showed slight greater variability across samples. The mean pairwise Jaccard similarity was 0.854 and the mean Dice similarity was 0.910, indicating generally strong agreement between runs. However, the mean terms ratio was lower than in the mitochondrial dataset, equal to 0.573. Although many extracted phenotypes are reproducible, OLIDA contains a larger fraction of HPO terms that appeared only intermittently across multiple executions. The results suggest that PhenoXtract is reliable in recovering a stable core of HPO terms.

Reproducibility metrics can provide complementary information to performance measures and may be incorporated as an additional evaluation criterion for LLM-based extraction tools.

### S.2. Additional Figures

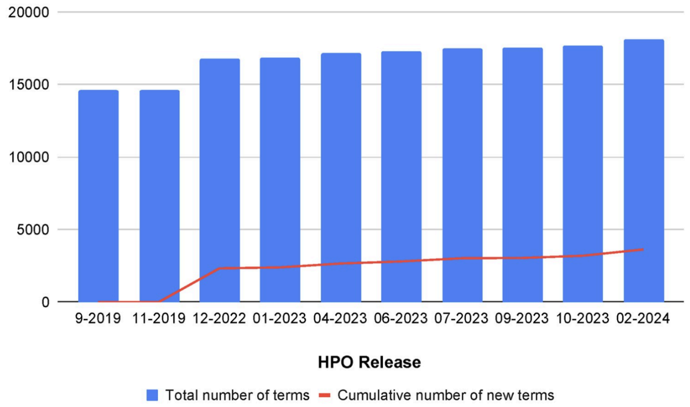


**Fig. S1:** HPO terms evolution. The evolution of the size of HPO in terms of number of concepts between December 2022 and February 2024. The line in the chart represents the growth in new terms being added to the ontology, which for this period was of 1303 terms, i.e. 7% increase. Source: [9].

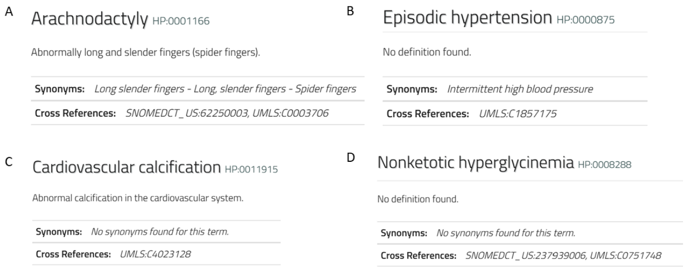


**Fig. S2:** Examples of HPO terms requiring preprocessing. A. No preprocessing needed; B. No definition: preprocessing to retrieve it. C. No synonyms: preprocessing to retrieve it. D. Neither synonyms nor definition: preprocessing to retrieve both.

A

At birth the child presented with macrocephaly, a wide anterior fontanelle, hypertelorism, polydactyly of hands and feet, hypogenitalism, cryptorchidism and congenital dislocation of the hips. At age 6 years he was severely retarded, without speech, markedly hypotonic and unable to stand or walk. His head circumference, and inner- and outer canthal distances were above the 97th percentile. Dysmorphic signs included downslanting palpebral fissures; short philtrum with upturned upper lip; open mouth with normal palate; normally shaped and posteriorly rotated ears and a broad and prominent forehead.

B

Regarding the onset, five of the patients were pediatric patients. Two were APAH-CHD patients, and three were diagnosed as IPAH patients. These three pediatric idiopathic patients had coincidental defects; patient three (HTP973) and patient six (MSD) had a patent foramen ovale, and patient seven (HTP1031) had a patent foramen ovale and a small patent ductus arteriosus. Regarding the two adult patients, one was associated with connective tissue disease (patient HTP964) and the remaining one was diagnosed with IPAH.

**Fig. S3:** Examples of clinical descriptions. A: Acrocallosal syndrome (ACLS) patient phenotypes description [10]. B: Phenotypes of seven patients with SOX17 Related Pulmonary Arterial Hypertension [11].

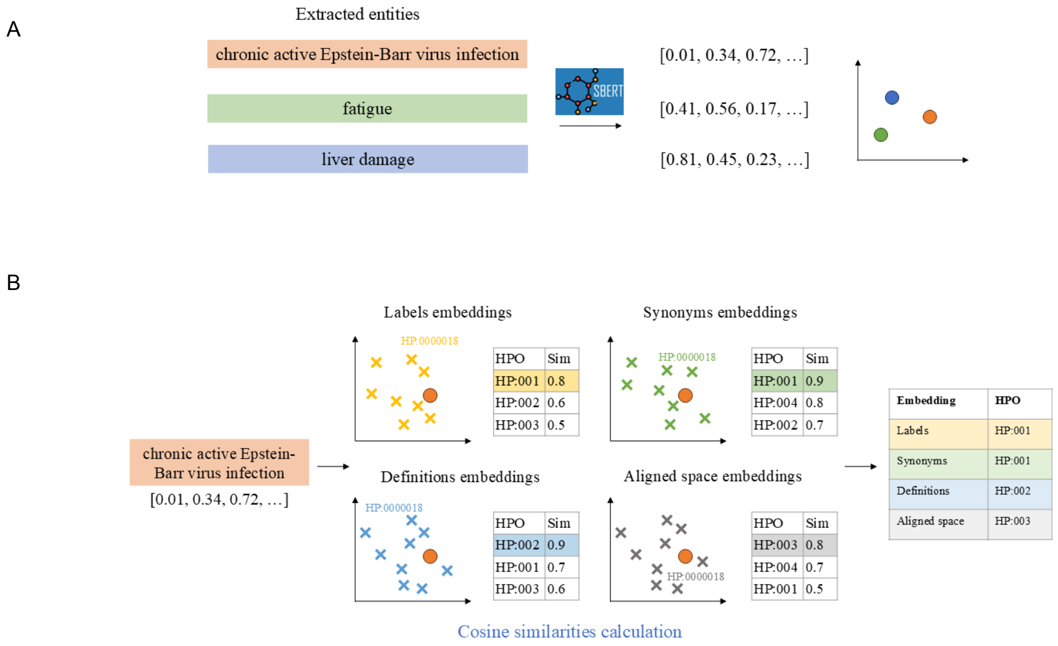


**Fig. S4:** Entity linking workflow. (A) generation of embeddings for entities extracted during concept recognition; (B) cosine similarity-based matching against ontology embeddings. A: Embeddings of CR extracted entities. Example of the first EL step: each entity extracted during CR is embedded using a Sentence Transformer (model: *all-MiniLM-L6-v2*), resulting in a numerical vector representation. B: Entity linking cosine similarity computation. Each entity embedding is compared against ontology term embeddings using cosine similarity, and the top-ranked phenotype per category is retained.

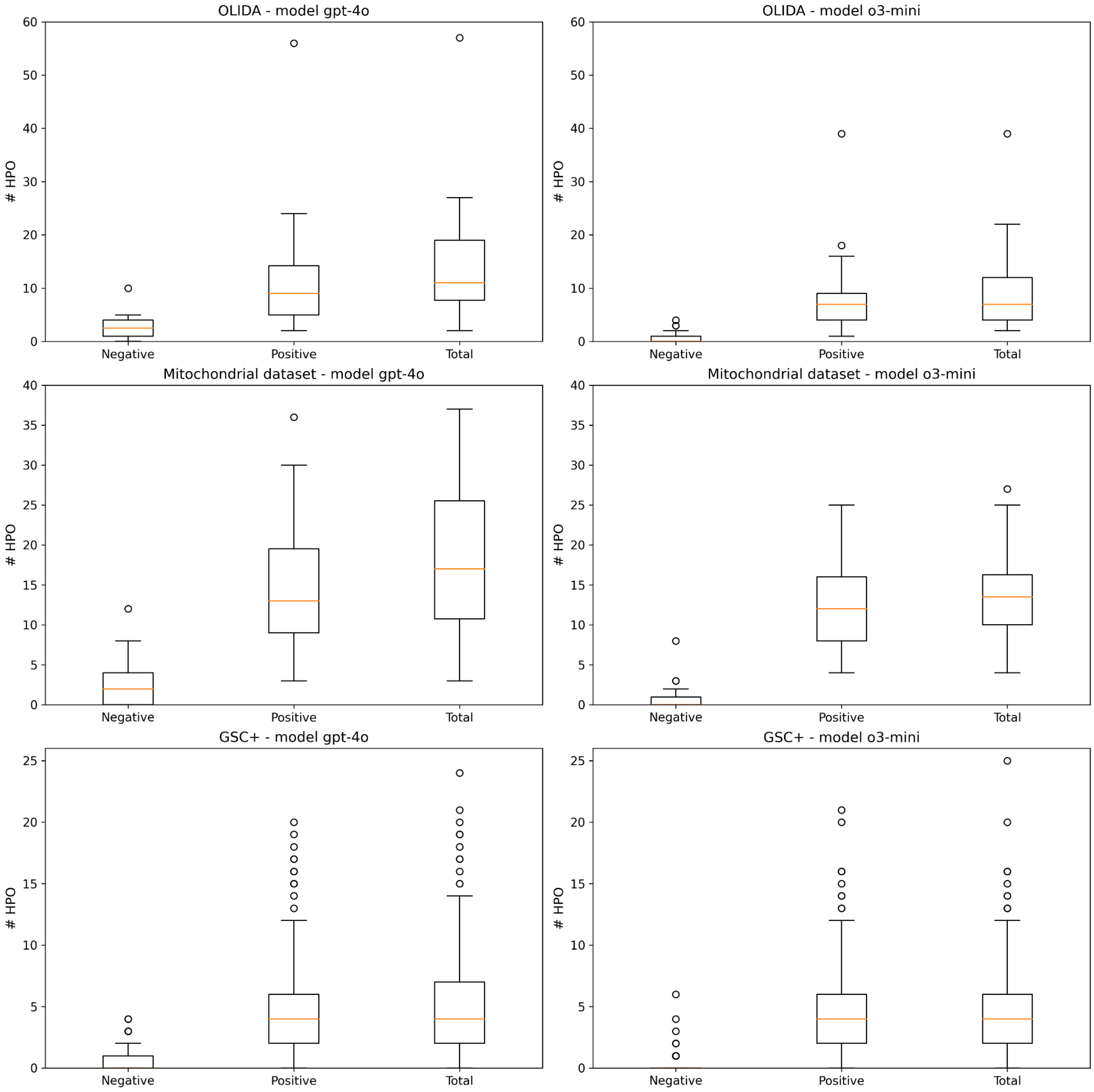


**Fig. S5:** Boxplots for each ground-truth of the number of positive HPO and negative HPO extracted from each input paper, produced with both model *gpt-4o* and model *o3-mini*. OLIDA, mitochondrial and GSC+ ground-truths have a median number of 9 HPOs (mean=11, total=259), 11 HPOs (mean=13, total=496) and 8 HPOs (mean=9, total=1933) per article, respectively.

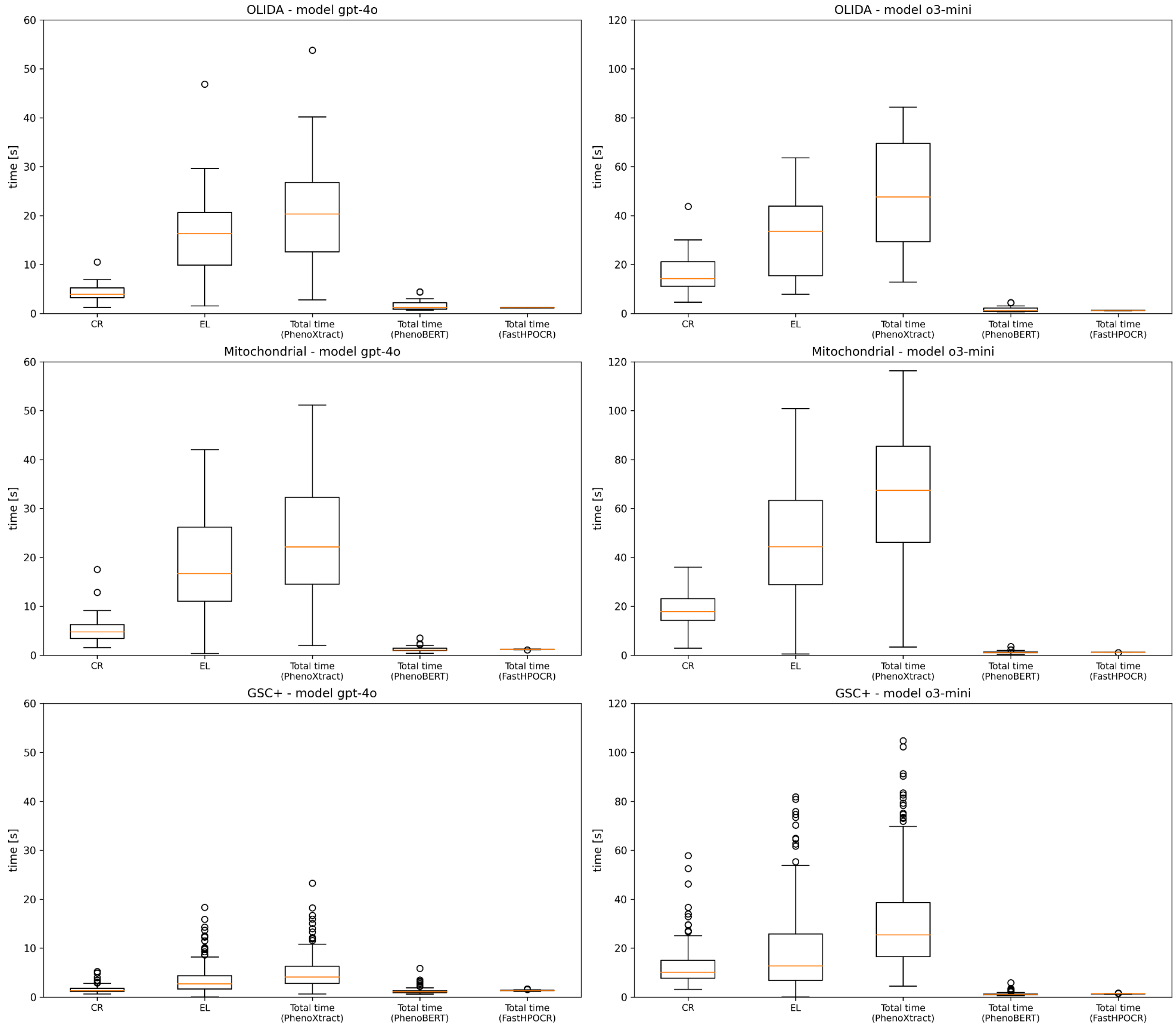


**Fig. S6:** PhenoXtract timing statistics. Timing distribution (Boxplots) for each dataset and model for CR, EL and the entire process (Total time = CR + EL).

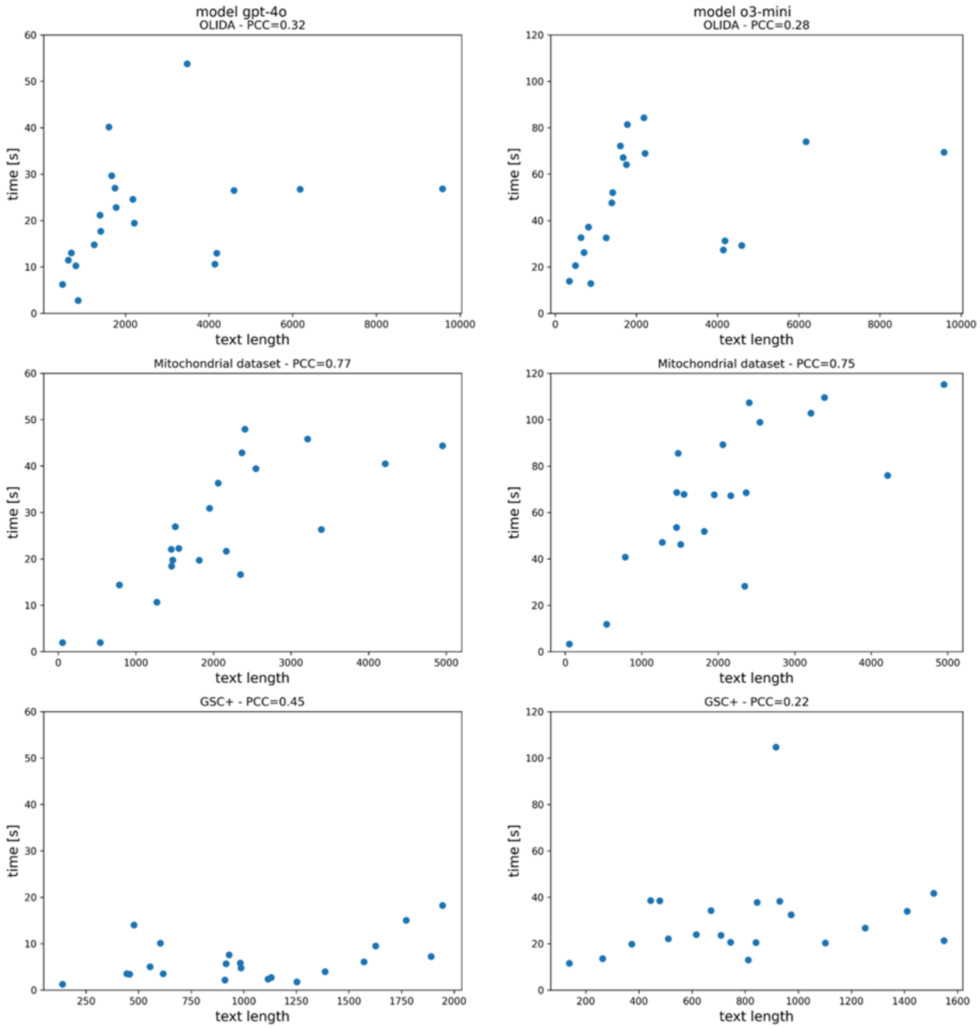


**Fig. S7:** Correlations between total execution time and input text length. Correlation between total execution time and input text length for OLIDA, mitochondrial dataset and GSC+ with model *gpt-4o* and model *o3-mini*. To investigate the characteristics most involved in execution time, correlations between the length of the analyzed text and the total time were plotted. Pearson correlation coefficient is shown in the individual titles. OLIDA shows quite low PCC (PCC=0.32 with *gpt-4o* and PCC=0.28 with *o3-mini*). Mitochondrial dataset, instead has higher values of PCC (PCC=0.77 with *gpt-4o* and PCC=0.75 with *o3-mini*). Finally, GSC+ does not show elements of correlations (PCC=0.45 with *gpt-4o* and PCC=0.22 with *o3-mini*).

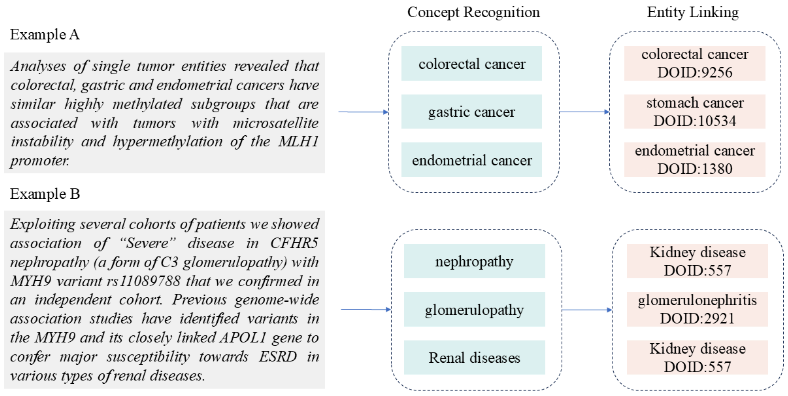


**Fig. S8:** Disease recognition and entity linking to DO. The two-step approach of PhenoXtract is applied to Disease Ontology. Concept Recognition (CR) and Entity linking (EL) results of two examples (Example A from [12] and B from[13]).

S.3. Additional Tables

| | |
|---|---|
| Base Model | microsoft/MiniLM-L12-H384-uncased |
| Max Sequence Length | 256 |
| Dimensions | 384 |
| Suitable Score Functions | dot-product, cosine-similarity, euclidean distance |
| Size | 120 MB |
| Pooling Mean | Pooling |
| Training Data | 1B+ training pairs |

**Table S4:** SentenceBERT *all-MiniLM-L6-v2.*

| **HPO element** | **Output matrix** | **Method** |
|---|---|---|
| Labels | labels_embeddings | SBERT *all-MiniLM-L6-v2* |
| Synonyms | synonyms_embeddings | SBERT *all-MiniLM-L6-v2* |
| Definition | definitions_embeddings | SBERT *all-MiniLM-L6-v2* |
| Knowledge graph | graph_embeddings | PyG GraphSAGE |

**Table S5:** Embedding matrices with the specification on method exploited.

| Case | Rule | Final HPO term |
|---|---|---|
| 1) at least 3 top ranked terms are the same on the most likely term | majority consensus | The majority term |
| 2) In one of the 4 categories, a term has cosine similarity equal to 1.0 | perfect match | The term with cosine similarity of 1.0 |
| 3) disagreement among 4 top-ranked embeddings | disambiguation step | Final match (mapping or discard) with additional LLM |

**Table S6:** Decision rules for final linking The disambiguation step to solve case 3) utilizes an additional layer of LLM. The four top-high-ranked candidate entities are provided at a prompt, explicitly instructed to select the most contextually coherent and clinically appropriate term to link to the input entity. The disambiguation step employed the same LLM as the CR step, using either *gpt-4o* or *o3-mini* depending on the configuration.

| OLIDA ID | Disease | Reference (PMID) |
|---|---|---|
| OLI167 | Porphyria | 16390615 |
| OLI832 | Acrocallosal syndrome | 23142271 |
| OLI367 | Kallmann syndrome | 23643382 |
| OLI672 | Hypertrophic cardiomyopathy | 28223422 |
| OLI549 | Familial long QT syndrome | 28749435 |
| OLI268 | Hypertrophic cardiomyopathy; Arrhythmogenic right ventricular cardiomyopathy | 30716529 |
| OLI025 | Primary hemophagocytic lymphohistiocytosis | 30782130 |
| OLI555 | Normosmic congenital hypogonadotropic hypogonadism | 31042289 |
| OLI647 | Sudden infant death syndrome | 31970460 |
| OLI1215 | Heterozygous familial hypercholesterolemia | 33642439 |
| OLI1146 | Bardet-Biedl syndrome | 34691137 |
| OLI1101 | Rare genetic deafness | 34946889 |
| OLI132 | MODY; Normosmic congenital hypogonadotropic hypogonadism | 32171037 |
| OLI981 | Torsade-de-pointes syndrome with short coupling interval | 33664309 |
| OLI1214 | Normosmic congenital hypogonadotropic hypogonadism | 34539727 |
| OLI101 | Usher syndrome type 1 | 21569298 |
| OLI283 | Amyotrophic lateral sclerosis | 25382069 |
| OLI359 | Nephronophthisis | 28392475 |
| OLI463 | Hypobetalipoproteinemia | 29540175 |
| OLI031 | Congenital hypothyroidism | 30375286 |

**Table S7:** OLIDA subset: 20 OLIDA IDs (with relative disease and reference to publication) composing the benchmark used to evaluate PhenoXtract performances.

| Dataset-Model | Step | Median [s] | Mean [s] | SD [s] |
|---|---|---|---|---|
| OLIDA - *gpt-4o* | Concept Recognition | 3.95 | 4.34 | 2.18 |
| OLIDA - *gpt-4o* | Entity Linking | 16.36 | 16.59 | 10.24 |
| OLIDA - *gpt-4o* | Total Time | 20.31 | 20.93 | 11.94 |
| Mitochondrial - *gpt-4o* | Concept Recognition | 4.76 | 5.35 | 3.18 |
| Mitochondrial - *gpt-4o* | Entity Linking | 16.68 | 18.69 | 12.08 |
| Mitochondrial - *gpt-4o* | Total Time | 22.11 | 24.04 | 14.38 |
| GSC+ - *gpt-4o* | Concept Recognition | 1.34 | 1.53 | 0.63 |
| GSC+ - *gpt-4o* | Entity Linking | 2.67 | 3.52 | 3.05 |
| GSC+ - *gpt-4o* | Total Time | 4.05 | 5.05 | 3.44 |
| OLIDA - *o3-mini* | Concept Recognition | 14.20 | 17.03 | 9.24 |
| OLIDA - *o3-mini* | Entity Linking | 33.50 | 36.67 | 32.52 |
| OLIDA - *o3-mini* | Total Time | 47.68 | 53.70 | 37.31 |
| Mitochondrial - *o3-mini* | Concept Recognition | 17.81 | 18.17 | 7.77 |
| Mitochondrial - *o3-mini* | Entity Linking | 44.34 | 43.93 | 26.66 |
| Mitochondrial - *o3-mini* | Total Time | 67.44 | 62.10 | 32.03 |
| GSC+ - *o3-mini* | Concept Recognition | 10.20 | 12.25 | 7.58 |
| GSC+ - *o3-mini* | Entity Linking | 12.71 | 18.66 | 16.83 |
| GSC+ - *o3-mini* | Total Time | 25.36 | 30.91 | 19.94 |

**Table S8:** Timing statistics (median, mean, and standard deviation) for each dataset and model. With the model gpt-4 OLIDA (20.93 ± 11.94 seconds, mean ± SD), compared to Mitochondrial dataset (24.04 ± 14.38 seconds), has lower total execution times. Likewise, with the model o3-mini, OLIDA (53.70 ± 37.31 seconds) has lower total execution times compared to Mitochondrial dataset (62.10 ± 32.03 seconds). Median time for model gpt-4 (OLIDA: 20.31 seconds, Mitochondrial dataset: 22.11 seconds, GSC+: 25.36 seconds) is less than the half of the median time for model o3-mini (OLIDA: 47.68, Mitochondrial dataset: 67.44, GSC+: 4.05).
